# Influences of chemotype and parental genotype on metabolic fingerprints of tansy plants uncovered by predictive metabolomics

**DOI:** 10.1101/2023.02.16.528607

**Authors:** Thomas Dussarrat, Rabea Schweiger, Dominik Ziaja, Thuan T. N. Nguyen, Liv Krause, Ruth Jakobs, Elisabeth J. Eilers, Caroline Müller

## Abstract

Intraspecific plant chemodiversity shapes plant-environment interactions. Within species, chemotypes can be defined according to variation in dominant specialised metabolites belonging to certain classes. Different ecological functions could be assigned to these distinct chemotypes. However, the roles of other metabolic variations and the parental genotype of the chemotypes remain poorly explored. Here, we first compared the capacity of terpenoid profiles and metabolic fingerprints to distinguish five chemotypes of common tansy (*Tanacetum vulgare*) and depict satellite metabolic differences. Metabolic fingerprints captured higher satellite variation while preserving the ability to define chemotypes. These satellite differences might influence plant performance and interactions with the environment. Next, to characterise the influence of the maternal genotype on chemodiversity, we performed variation partitioning and generalised linear modelling. Our findings revealed that maternal genotype was a higher source of chemical variation than chemotype. Predictive metabolomics unveiled 184 markers predicting maternal genotype with 89% accuracy. These markers included, among others, phenolics, whose functions in plant-environment interactions are well established. Hence, these findings place parental genotype at the forefront of intraspecific chemodiversity. We thus recommend considering this factor when comparing the ecology of various chemotypes. Besides, the combined inclusion of inherited and satellite metabolic variation in computational models may help connecting chemodiversity and evolutionary principles.

## Introduction

The evolution of plant metabolism has attracted the curiosity of scientists for decades^1–3^. Sources of chemical diversity in plants are abundant, while explanations of its role and origin are not fully elucidated, especially regarding intraspecific chemodiversity^4,5^. Yet, previous studies revealed high intraspecific chemodiversity in various species and linked metabolic variation to considerable ecological consequences^6–8^. In some plant species expressing high intraspecific chemodiversity, individuals can be classified into distinct chemotypes according to the occurrence and ratio of individual compounds belonging to a specific major metabolite class such as terpenoids for aromatic plants, glucosinolates for Brassicaceae or pyrrolizidine alkaloids for Asteraceae^9–11^. The strategy of discriminating plants according to their chemotypes can reveal interesting information and improve our comprehension of the ecology and evolution of intraspecific chemodiversity^10–12^. For instance, slugs show distinct preferences for certain *Solanum dulcamara* (Solanaceae) chemotypes, which are determined by the glycoalkaloid composition^13^. Evolution in the cardenolide profile could be linked to the surrounding biotic pressure and confer various toxic properties in *Erysimum* (Brassicaceae) species^14^. Moreover, high intraspecific diversity in plant chemotypes may be crucial for invasion success and different chemotypes may hence show distinct geographic distributions^15,16^. However, these chemotypes may capture a significant fraction of intraspecific chemodiversity but do not fully cover the chemodiversity blend. In fact, other pivotal metabolites or metabolite classes, here called satellites, may differ between chemotypes and have distinct ecological functions^8,17^. The correlation between satellite metabolites and the main compounds that determine chemotypes has rarely been looked at, although the assumed restriction to the major chemical class to define chemotypes could be a source of misinterpretation.

Furthermore, chemotypes are heritable^10,18^, but the inheritance does not always follow Mendelian laws^19,20^. In addition, plants of a given chemotype can have distinct genetic backgrounds^19^. Thus, the parental genotype might also be responsible for a substantial part of chemodiversity. In this scenario, extending the research from main chemical patterns (defining chemotypes) to potential chemical variation inferred by parental genotype may considerably increase our understanding of intraspecific chemodiversity. This approach may also contribute to deciphering the genetic laws governing chemodiversity inheritance. Thus, large scale metabolomics analyses of highly chemodiverse species may help to characterise the nature of satellite metabolic diversity within chemotypes and explore the impacts of the parental genotype on the metabolic variation.

Common tansy [*Tanacetum vulgare* L., also known as *Chrysanthemum vulgare* (L.) Bernh. Asteraceae], possesses an astonishing intraspecific chemodiversity^8^. Tansy chemotypes are defined by their dominant terpenoid(s), which can contribute 41-99% of the leaf total terpenoid profile. Mono-chemotypes contain one dominant terpenoid (more than 50%) and commonly found examples in the field are the β-thujone chemotype or the camphor chemotype^1,6,8^. The mixed-chemotypes comprise two to three dominant terpenoids. Next to these dominant terpenoids, tens of further terpenoids can be found in both mono- and mixed chemotypes, contributing to the full terpenoid bouquet. These different chemotypes can co-occur and up to 14 distinct chemotypes were found in a rural area of just a few square kilometers^6^. Previous studies highlighted the consequences of this intraspecific chemodiversity on insect behaviour and performance as well as chemotype-specific differences in chemical responses to herbivory and abiotic constraints^12,21,22^. Offspring individuals of one mother plant vary in terpenoid profiles and other chemical classes since tansy is outcrossing^19,23^. Hence, tansy represents an ideal study system to test the nature of satellite variation within chemotypes and investigate whether the parental genotype influences intraspecific chemodiversity.

To meet these objectives, terpenoid analyses using gas chromatography-mass spectrometry (GC-MS) as well as untargeted metabolic fingerprinting using ultra high performance liquid chromatography coupled to a quadrupole time-of-flight-MS (UHPLC-QTOF-MS/MS) were performed on leaves collected from five chemotypes of tansy plants that had been grown in a semi-field experiment. The chemotypes were derived from different maternal genotypes. Generalised linear models (GLMs) were conducted to test (i) whether metabolic features detected by untargeted metabolic fingerprinting were as performant as terpenoid profiles to predict chemotypes and (ii) whether the maternal genotype significantly affected the metabolome of tansy. The predictive capacity of certain metabolic markers was validated by resampling plants in the field and clones of these plants grown in the greenhouse. Potential consequences of our findings on the interpretation of (chemo-) ecological experiments are discussed.

## Results

### Tansy chemotypes are mainly defined by quantitative variation in their metabolome

As a first step to characterise the variation in satellite metabolites, we captured the leaf terpenoid profiles (GC-MS) and metabolic fingerprints (UHPLC-QTOF-MS/MS) of five tansy chemotypes (181 samples) obtained from fourteen maternal genotypes (four to eight per chemotype) and grown in a semi-field common garden (for analytical workflow see Fig. 1). The chemotypes included two mono-chemotypes, artemisia ketone (in the following called “Keto”) and *β*-thujone (“BThu”), and three mixed-chemotypes, dominated by either *α*-thujone and *β*-thujone (“ABThu”), or artemisyl acetate, artemisia ketone and artemisia alcohol (“Aacet”) or (*Z*)-myroxide, santolina triene and artemisyl acetate (“Myrox”). In the field, plants were either grown in homogenous plots, consisting of five plants of the same chemotype, or in heterogenous plots, consisting of plants of five distinct chemotypes. Leaves without visible damage were sampled for chemical analyses in June 2021. In total, 52 compounds (mostly terpenoids) were detected by GC-MS, while untargeted LC-MS analyses yielded 5,066 features (Tables S1 & S2). Growth conditions (*i.e.* plants grown in homogenous or heterogenous plots) did not show an effect on the terpenoid profiles (Fig. S1). In contrast, the five chemotypes were distinguishable by a PCA and displayed interesting chemical patterns based either on terpenoid profiles or LC-MS features (Fig. 2). The major discriminant terpenoids were congruent with the chemotype definition^6,19^ and included *α-* and *β*-thujone, artemisia ketone, artemisyl acetate and artemisia alcohol, and santolina triene and (*Z*)-myroxide.

**Fig. 1.**
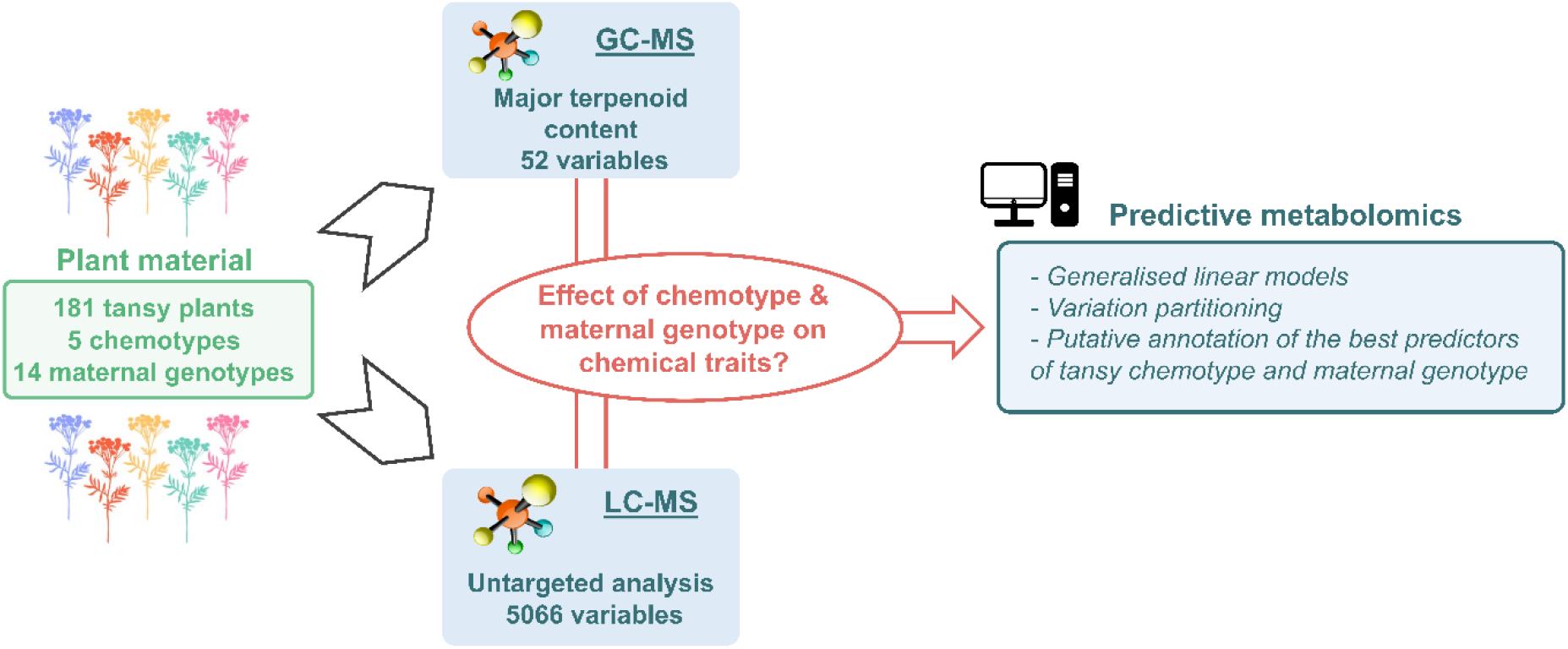
Simplified scheme of the analytical workflow.

**Fig. 2.**
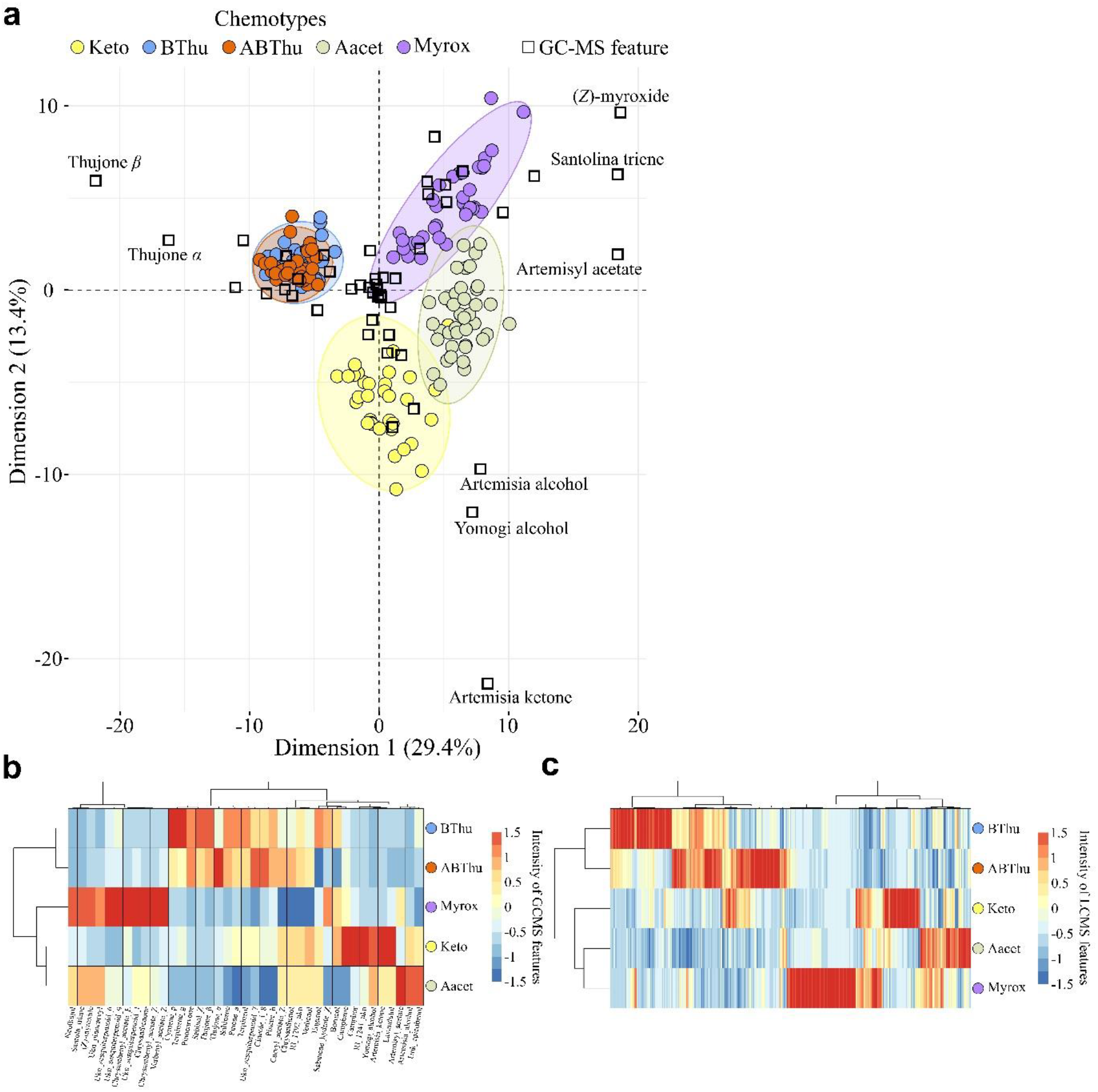
Identification of tansy chemotypes via GC-MS and LC-MS analyses. **a.** Principal component analysis biplot illustrating the discrimination of tansy chemotypes using GC-MS data (52 features). **b.** Representation of the levels of the 39 significant GC-MS features (Tukey’s test, *P* < 0.05, FDR correction) differentiating chemotypes via clustering analysis (Pearson correlation, Ward algorithm). **c.** Clustering analysis (Pearson correlation, Ward algorithm) of tansy chemotypes using 809 significant LCMS features (Tukey’s test, *P* < 0.05 after FDR correction). *Keto:* artemisia ketone chemotype, *Bthu:* β-thujone chemotype, *ABThu:* α-/β-thujone chemotype, *Aacet*: artemisyl acetate/artemisia ketone/artemisia alcohol chemotype, *Myrox*: (*Z*)-myroxide/santolina triene/artemisyl acetate chemotype.

Next, we questioned whether this discriminatory ability was due to qualitative and/or quantitative variation. Notably, 79% of the features detected using both GC-MS and LC-MS occurred in all five chemotypes and 97% of the features were detected in more than one chemotype (Table S3). A rather high quantitative variation was observed, with 39 GC-MS and 809 LC-MS features differing significantly in abundance among chemotypes (Tukey’s test, *P* < 0.05, FDR correction) (Fig. 2b and 2c). Altogether, these results suggested that the distinction between tansy chemotypes more likely resulted from quantitative variation rather than from differences in the occurrence of specific compounds.

### Major terpenoid profiles and metabolic fingerprints show comparable chemotype predictive performance

To compare the predictive capacity of either terpenoid profile (*i.e.* GC-MS) or metabolic fingerprint (*i.e.* LC-MS), we deployed generalised linear models (GLMs) dividing the sample set into a “training” set, a “validation” set and a “test” set. Chemotypes were considerably predictable (Fig. 3a). Significant GCs-MS or LC-MS features resulted in an average accuracy of 95% and 93%, respectively. The selection of the most predictive features (here called best markers/predictors) was carried out based on their occurrence in the 500 models. The predictive performance of the 39 best LC-MS predictors was then compared to that of the 39 GC-MS predictors (significant terpenoids) (Tab. S4). Subsequently, the seven terpenoids used to define the five tansy chemotypes chosen for our study were tested (*i.e. α-* and *β*-thujone, artemisia alcohol, artemisia ketone, artemisyl acetate, (*Z*)-myroxide and santolina triene). Chemotypes were predicted at 95%, 98% and 97% accuracy using the 39 significant terpenoids, the 7 chemotype-defining terpenoids or the best 39 LC-MS markers, respectively (Fig. 3a). The GLM-based modelling approach was statistically validated by evaluating the likelihood of spurious prediction using 500 permuted datasets where chemotypes were randomly permuted between samples, yielding a 5% accuracy. To test the robustness of the LC-MS markers, we used a complementary dataset composed of field and greenhouse plants collected in October 2022. Thirty-four of the 39 features were refound in this new dataset based on their retention time, *m/z* ratio and MS/MS spectra. Model equations were defined on samples collected and analysed in June 2021 and directly applied to the samples taken in 2022 to predict their chemotype. The average prediction accuracy of 90% demonstrated the robustness of the markers across years, seasons and growing conditions (Fig. 3a).

**Fig. 3.**
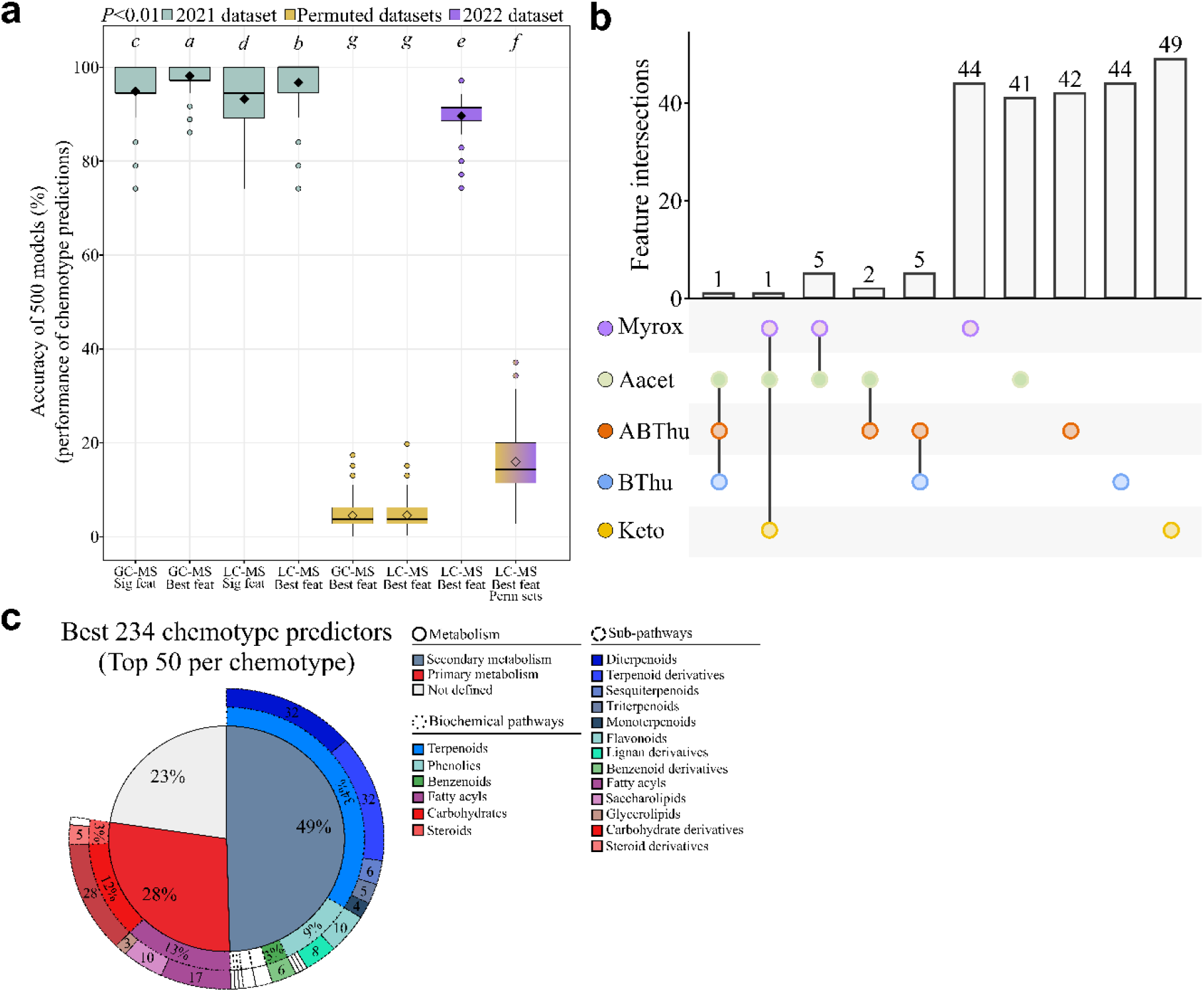
Predictive metabolomics on tansy plants. **a.** *R^2^* scores of the 500 generalised linear models using GC-MS or LC-MS features (Tukey’s test, *P* < 0.01). For each condition, 500 permuted datasets were created by randomly swapping chemotype classes. *Feat:* features, *Perm:* permuted. *Sig:* significant. “GC-MS Sig feat” included 39 significant features, “GC-MS Best feat” contained 7 features, “LC-MS Sig feat” contained 809 significant features, “LC-MS 39 feat” included 39 features. **b.** Upset plot of the top 50 markers for each chemotype from LC-MS modelling. The absence of a dot means that the corresponding markers were not present among the best 50 markers in the corresponding chemotypes. **c.** Putative annotation of the best LC-MS predictors. *Keto:* artemisia ketone chemotype, *BThu:* β-thujone chemotype, *ABThu:* α-/β-thujone chemotype, *Aacet:* artemisyl acetate/artemisia ketone/artemisia alcohol chemotype, *Myrox:* (*Z*)-myroxide/santolina triene/artemisyl acetate chemotype.

Next, we analysed the intersections between the top 50 markers for each chemotype (Tab. S4). The best predictors strongly differed among chemotypes (Fig. 3b). However, their status of serving as best markers did not lie in their exclusivity but rather in variation in their abundance, since only 11% of the markers were specific to one chemotype (Fig. S2 & Tab. S3). We then explored the relationships between the best 39 LC-MS markers and the 7 terpenoids used to define our five chemotypes, yielding strong correlations (Fig. S3). Overall, these results demonstrated that both terpenoid profiles and metabolic fingerprints could be used to predict tansy chemotypes efficiently, thus questioning the chemical nature of the best predictive LC-MS features.

### Major terpenoids as the main predictors of tansy chemotypes, which are further defined by satellite metabolic variation

To gain further insight into the biochemical pathways characterising tansy chemotypes, we putatively annotated the top 50 LC-MS markers per chemotype using MS and MS/MS spectra. The MSI annotation level for each marker is presented in Tab. S5. The majority of the best predictors were putatively assigned as specialised (secondary) metabolites (Fig. 3c). As expected, terpenoids were overrepresented among the markers and included putative diterpenoids and terpenoid derivatives (32 compounds each), as well as sesqui-, tri- and monoterpenoids. Further markers were putatively assigned to other biochemical pathways such as phenolics (18 compounds, 9% of the best markers). Primary metabolites such as fatty acyls and carbohydrate derivatives were also found (Fig. 3c). Overall, while this analysis supported the central place of major terpenoids to define chemotypes of tansy, findings highlighted variation in satellite metabolites within chemotypes that should be considered in further studies.

### Predictive metabolomics sheds light on the influence of maternal genotype on metabolic fingerprints

While chemotypes could be used to classify tansy samples efficiently, the parental genotype may also significantly impact intraspecific chemodiversity. To test this hypothesis, we performed variation partitioning (Fig. 1). Chemical variation in tansy metabolism was influenced by both chemotype and maternal genotype using either GC-MS or LC-MS data (Fig. 4a and 4b). The terpenoid profile was mainly determined by the chemotype (64.7%) and only 3.2% of the variation was exclusively explained by the maternal genotype. Conversely, the maternal genotype explained 17.3% of the chemical variation in metabolic fingerprints independently of the chemotype. Chemotype explained 1.4% of the variation. In total, 41 (out of 52) terpenoids and 3,688 (out of 5,066) LC-MS features displayed significant differences among the 14 maternal genotypes (Tukey’s tests, *P* < 0.05, FDR correction) (Fig. S4). To further characterise the impact of maternal genotype on tansy metabolism, we employed GLMs. First, a clustering analysis was used to classify the 14 maternal genotypes into four main classes to allow GLM analyses by increasing the number of samples per class (Fig. S4). While GC-MS features could hardly predict maternal genotypes, the top 5% predictive LC-MS features (*i.e.* 184 features) predicted the maternal genotype with 89% accuracy. Models were statistically validated using 500 permuted datasets (Fig. 4c). Besides, the predictive performance of the best markers was biologically validated using a complementary set. Model equations were defined on samples collected in June 2021 and directly applied to the complementary set composed of 20 plants harvested in the field in October 2022, yielding an average accuracy of 69% (Fig. 4c). While this result underlined the predictive capacity of the best markers across years and seasons, the 20% delta between 2021 and 2022 predictions could be explained by (i) a low number of samples per class in the complementary set, which increased error weight on tested samples and/or (ii) a slightly different abundance of these markers between seasons. Predictions in permutation tests (accuracy of 30%) were ascribed to the low number of samples per maternal genotype class in the complementary set. Thus, these results highlighted the strong influence of maternal genotype on the tansy metabolome. Besides, only 33% of the 184 LC-MS markers, which predicted maternal genotype with 89% accuracy, also differed significantly in abundance among chemotypes (Tab. S5), supporting the hypothesis that chemotypes only capture part of the entire intraspecific chemodiversity.

**Fig. 4.**
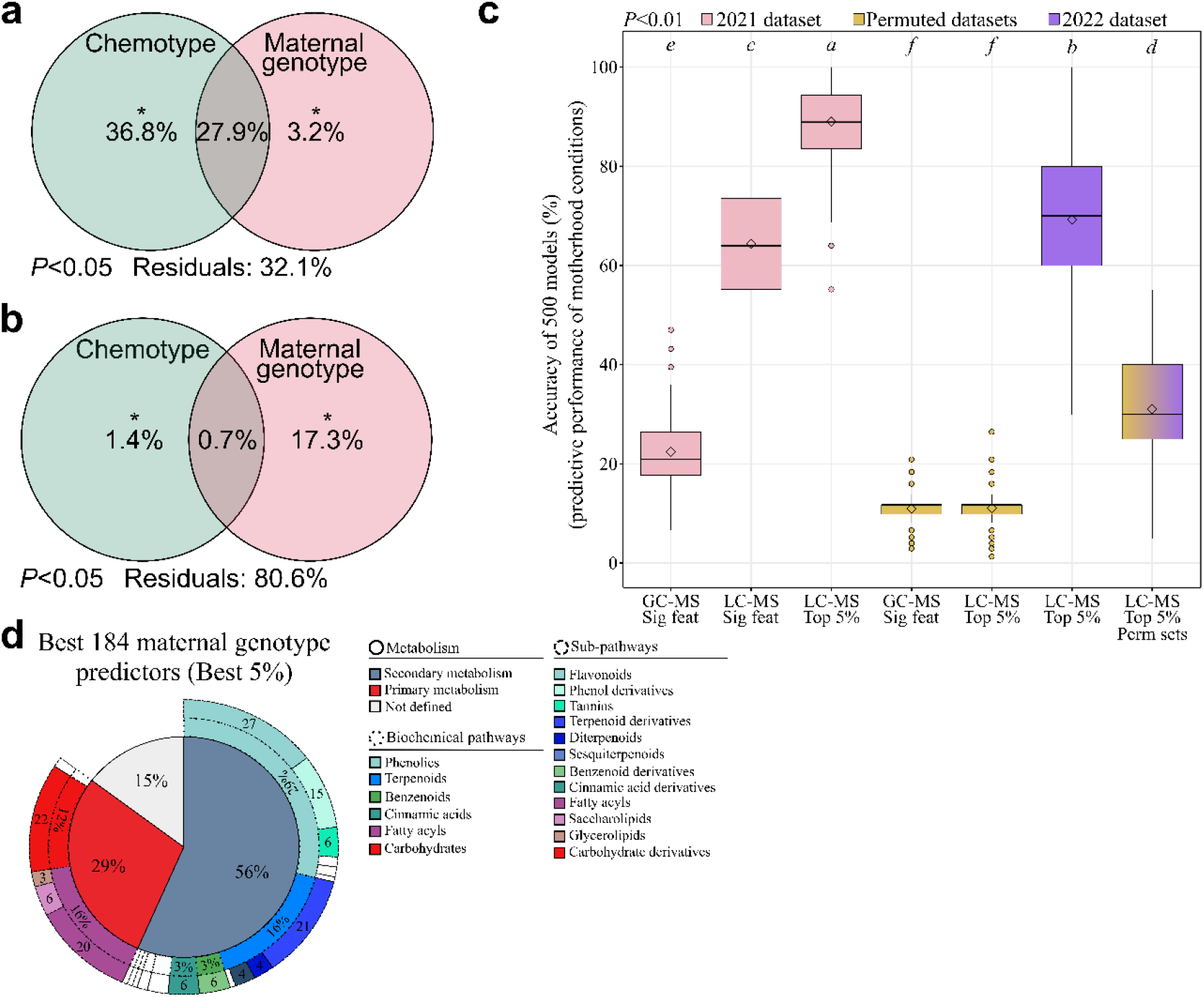
Effect of maternal genotype on intraspecific chemodiversity. **a.** Variation partitioning on 52 GC-MS features (ANOVA test, *P* < 0.05). **b.** Variation partitioning on 5,066 LC-MS features (ANOVA test, *P* < 0.05). **c.** *R2* scores of the 500 generalised linear models using GC-MS or LC-MS features (Tukey’s test, *P* < 0.01). For each condition, 500 permuted datasets were developed. *Feat:* features, *Perm:* permuted. *Sig*: significant. “GC-MS Sig feat” included 41 significant features, “LC-MS Sig feat” contained 3,688 significant features and “LC-MS Top 5%” included the best 5% predictors (184). **d.** Putative annotation of the best LC-MS predictors.

To test the influence of maternal genotype on the tansy metabolome within chemotypes, we deployed partial least square discriminant analyses. While GC-MS data only slightly discriminated maternal genotypes within chemotypes, a clear distinction was found using LC-MS data (Figs S5, S6). Hence, maternal genotype significantly influenced the overall intraspecific chemodiversity and represented a significant source of chemical variation within chemotypes.

### Chemical variation in primary and specialised metabolism among maternal genotypes

To define the main metabolic pathways impacted by maternal genotype, we putatively annotated the best 184 LC-MS markers (*i.e.* top 5%) (Tables S5, S6). Maternal genotype influenced both primary and specialised metabolism (Fig. 4d). Phenolics were the most affected class (29%), including flavonoids (27 compounds) such as quercetin and derivatives of quercetin, kaempferol, luteolin and naringenin (Table S5). To a lower extent, terpenoids, fatty acyls and carbohydrate derivatives were also impacted. Moreover, cinnamic acid derivatives, benzenoids and nitrogen-related compounds were represented among the best markers (Fig. 4d, Table S5). Thus, these findings highlighted the strong metabolic variation induced by the maternal genotype, including variation in major classes such as flavonoids, which could in turn lead to significant ecological consequences.

## Discussion

### Untargeted metabolomics as a complementary strategy to study intraspecific chemodiversity

The analysis of intraspecific chemodiversity offers promising perspectives to improve our comprehension of the evolution of plant specialised metabolism and ascribe ecological functions to specific chemical traits^4^. In this context, highly chemodiverse species are often classified into chemotypes based on prominent and dominant specialised metabolites, such as terpenoids in tansy^6^. Terpenoid chemotypes have also been described in numerous weed species, such as *Solidago gigantea* (Asteraceae) and plants used as spices or for medical purposes, such as *Thymus vulgaris* (Lamiaceae) or *Cannabis sativa* (Cannabaceae)^24–26^. These examples highlight the fascinating chemical polymorphism that can be found even within species. Different chemotypes may show differential invasion success^25,27^ and are exposed to differential selection by various herbivores, such as aphids or slugs^13,28^. Thus, the classification in chemotypes can offer highly valuable insights. However, this strategy restricts our comprehension of chemotype consequences for ecological interactions to a specific compound class^8^. Strikingly, our untargeted analysis showed that changes in the main terpenoid profiles were accompanied by a significant variation in numerous satellite metabolites, which may also be of ecological importance. GLMs displayed a comparable performance of GC-MS and LC-MS data to predict chemotypes, thus placing LC-MS analysis as a valuable alternative to define chemotypes. Nevertheless, compared to LC-MS analyses, which may be supplemented by measurements in positive electrospray ionisation mode, GC-MS analyses represented a more efficient strategy to depict the terpenoid profile. Besides, the terpenoid profile remained the best predictor for chemotypes. However, in contrast to GC-MS, LC-MS measurements captured the importance of variation in satellite metabolites in defining chemotypes. Prominent satellite chemical variation in the tansy chemotypes was found, for example in phenolics, which was consistent with the chemodiversity recently described in this compound class among *Chrysanthemum* species^29^. Additional variation was observed in fatty acyls and carbohydrate derivatives, which is congruent with previous studies on tansy^8,30^.

Analysing variation in satellite metabolites may be of relevance for various objectives in the research area of intraspecific chemodiversity. First, these satellite metabolites need to be taken into consideration when ascribing functional roles to a certain chemotype, as functions should not be assigned based on the dominant terpenoid(s) only^8^. For instance, phenolics also showed protective functions against aphids^31^. Second, satellite chemical variation may be of importance when exploring intra-individual chemodiversity among organs or tissues. For example, while tansy chemotypes are usually discriminated based on their leaf terpenoids, flowers may show slightly distinct terpenoid patterns and have been found to explain preference and abundance patterns of pollinators and florivores^12,32^. Other metabolite classes of flower parts, such as protein and lipid contents of pollen, did not necessarily differ among different tansy leaf chemotypes^12^. Besides, leaf chemotypes and phloem sap chemistry were only partially linked in tansy plants^22^ while root and leaf terpenoid profiles showed mostly uncorrelated patterns^33^. Hence, analysis of other metabolites or classes of metabolites apart from the major chemotype-determining terpenoids clearly benefits our knowledge of chemodiversity within plant individuals. Third, further investigation of satellite chemodiversity may help deciphering the genetic rules governing the inheritance of intraspecific chemodiversity. Thereby, the inheritance may differ for different biosynthetic pathways such as terpenoids and phenolics^34,35^. Finally, exploring the correlations between terpenoids and other specialised or also primary metabolites may support the development of computational models that aim to link chemodiversity and evolutionary principles^36,37^.

### Impacts of maternal genotype on intraspecific chemodiversity and potential ecological consequences

Variation partitioning indicated that variation in tansy metabolism was not exclusively driven by chemotype but rather derived from maternal genotype. Chemical variation in the tansy metabolome was explained at 18% by maternal genotype and 184 markers predicted this parameter with 89% accuracy. In other words, the chemotype was responsible for only certain facets of intraspecific chemodiversity in tansy. The additional chemical variation inferred from the maternal genotype could have significant consequences. For instance, terpenoids, phenolics, benzenoids and fatty acyls were among the most represented metabolite classes within the best markers for maternal genotypes. The role of terpenoids in the attraction from the distance of herbivores, their natural enemies and pollinators is well established^38,39^. These compounds also served plant fitness by acting as repellents or defensive compounds against several antagonists^40,41^. Several other compounds can influence herbivore performance. For instance, phenolic glycosides displayed defensive functions against generalist herbivores^42^. Similar effects have been reported for certain cinnamic acid derivatives and tannins^43^. Besides, the occurrence of several flavonoids and other phenolics, such as quercetin, kaempferol, luteolin and naringenin derivatives as well as caffeoylquinic acids, is in agreement with previous reports on tansy biochemistry^44,45^. Flavonoids can also influence the behaviour of belowground organisms by either conferring stimulatory or deterrent properties^46–48^. In contrast, benzenoids have mostly been recognised for their role in pollinator attraction^49^. Furthermore, several metabolites found here as markers for maternal genotypes, such as terpenoids and fatty acyls, are known to affect plant responses to herbivory under challenging abiotic conditions^21,50^.

Moreover, the effect of the maternal genotype was not only observable at the metabolic fingerprint scale, but also within chemotypes of tansy. Hence, the maternal genotype represents at least partially the observed satellite metabolic variation within chemotypes. This observation can be supported by the fact that the reliability of genotype in predicting chemotype depends on the organism^51^. In addition, a given chemotype may arise from distinct parental genotypes. Since the inheritance of the chemotype in tansy and other species is assumed to be complex and defined by a combination of genes^19,20^, the concentrations of major compounds such as terpenoids can vary according to this genetic combination and thus be distinguished between parental genotypes within a given chemotype. Moreover, the transmission of genes allowing chemotype inheritance could be accompanied by additional genetic information governing other metabolic patterns, which could be distinguished within tansy chemotypes^8,19^.

## Conclusion

Overall, our predictive untargeted metabolomics approach highlighted that (i) variation in terpenoid contents within chemotypes was accompanied by significant satellite metabolic variation and (ii) maternal genotype was a stronger driver of intraspecific chemodiversity than chemotype. Multiple consequences of this additional metabolic variation thus need to be considered, as discussed above, sensitising researchers to consider parental genotype effects when working with highly chemodiverse species. Analysing the parental genotype effect on a wider range of chemotypes and assessing the ecological consequences of satellite metabolic variation are exciting perspectives for intraspecific chemodiversity research.

## Materials and Methods

### Plant stock

Seeds of tansy were collected in the surroundings of Bielefeld (Germany) from different maternal plants grown at a distance of at least 20 m, assuming thus that these are different maternal genotypes. Seeds were germinated and leaf terpenoid profiles were determined from offspring by GC-MS, as described below. From these offspring, plants of five chemotypes were selected for further experiments, including two mono-chemotypes, BThu and Keto, and three mixed-chemotypes, ABThu, Aacet and Myrox. These plants were derived from the seeds of in total 14 different maternal plants. Overall, each chemotype was derived from four up to eight different maternal plants, and from each maternal plant, we used two to three different chemotypes. Thus, we had 150 different chemo-genotypes, which were kept as “stock” in a greenhouse since the end of 2019.

### Field experiment and leaf harvest

A semi-field common garden experiment was established in 2020 (52°03’39.43’N, 8°49’46.66’E) (for details see Ziaja and Müller, 2022^52^). The field was divided into six blocks, each comprising ten plots of five plants (1 m between plots and 2 m between blocks). From each of the stock plants, we prepared two clones (total 300 plants) and planted one in a homogenous plot, consisting of five plants of the same chemotype, and the other in a heterogenous plot, consisting of five plants of five distinct chemotypes. There was no obvious impact of these distinct growth conditions on the leaf terpenoid profile (Fig. S1). Importantly, all plants within a plot were descendants of distinct maternal plants. In June 2021, the tip of the youngest fully-expanded leaf without any visible herbivory damage or pathogen infestation was collected from each plant, directly frozen in liquid nitrogen and stored at −80 °C until freeze-drying and chemical analyses. Due to a freezer incident, only 181 samples (of the 300 collected) could be used for this analysis. This sample set included at least 35 samples per chemotype and 9 samples per maternal genotype (Table S7). Additional samples were collected following the same protocol from plants of the field experiment as well as from the greenhouse stock in October 2022 and used as a validation set (then called complementary set) for metabolic fingerprinting, testing the robustness of the determined metabolic markers (see below).

### Metabolite extraction

For terpenoid analyses by GC-MS, 10 ± 2 mg dried samples were extracted in *n*-heptane (99%, Carl Roth, Karlsruhe, Germany) containing 1-bromodecane as an internal standard (97%, Sigma Aldrich, Taufkirchen, Germany). Samples were sonicated for 5 min, centrifuged and supernatants were used for GC-MS analyses^52^. For analyses of metabolic fingerprints by UHPLC-QTOF-MS/MS, dried leaf material was homogenised and 8 ± 2 mg samples were extracted in methanol 90% (v:v) containing hydrocortisone as internal standard (Sigma-Aldrich, Steinheim, Germany) by sonicating in an ice bath for 15 minutes. Supernatants were collected, centrifugated for 10 minutes and filtered using 0.2 μm filters (Phenomenex, Torrance, CA, USA) as previously described^53^.

### Metabolomics

For GC-MS (GC 2010 Plus - MS QP2020, Shimadzu, Kyoto, Japan), separation was performed using a semi-polar column (VF-5 MS, 30 m length, 0.25 mm ID, 10 m guard column, Varian, Lake Forest, USA). The GC injection port was kept at 240 °C. The GC temperature program started at 50 °C kept for 5 min, increased to 250 °C at a rate of 10 °C min^-1^ and further increased to 280 °C at a rate of 30 °C min^-1^, which was held for 3 min. Electron ionisation at 70 eV was applied. An alkane standard mix (C7-C40, Sigma Aldrich) was analysed with the same method to determine the retention indices of the terpenoid analytes^54^. Terpenoid identification was performed by comparing mass spectra and retention indices to chemical references and chemical databases as previously described^52^.

For UHPLC-QTOF-MS/MS (UHPLC: Dionex UltiMate 3000, Thermo Fisher Scientific, San José, CA, USA; QTOF: compact, Bruker Daltonics, Bremen, Germany), the separation was performed on a Kinetex XB-C18 column (150 × 2.1 mm, 1.7 μm, with guard column; Phenomenex) at 45 °C and a flow rate of 0.5 mL min^-1^ using a gradient from eluent A, *i.e.* Millipore-H2O with 0.1% formic acid (FA), to eluent B (acetonitrile with 0.1% FA): 2 to 30% B within 20 min, increase to 75% B within 9 min, followed by column cleaning and equilibration, as described in Schweiger et al. (2021)^53^. The QTOF was operated in negative electrospray ionisation (ESI) mode at a spectra rate of 6 Hz in the *m/z* (mass-to-charge) range of 50-1300. The settings for the MS mode were: end plate offset 500 V, capillary voltage 3,000 V, nebulizer (N2) pressure 3 bar, dry gas (N2; 275 °C) flow 12 L min^-1^, low mass 90 m/z, quadrupole ion energy 4 eV, collision energy 7 eV. The AutoMS/MS mode was used to obtain MS/MS spectra ramping the isolation width and collision energy along with increasing *m/z.* Additional MS/MS analyses of some samples were performed to target selected ions (*i.e.* best markers) using multiple reaction monitoring (MRM). A calibration solution with sodium formate was used for the recalibration of the *m/z* axis. For the samples collected in 2021, raw LC-MS data were processed via DataAnalysis (v. 4.4, Bruker Daltonics) using optimised parameters, which included signal-to-noise ratio 3, correlation coefficient threshold 0.75, minimum compound length 20, smoothing width 3. Bucketing was applied to sort features belonging to the same metabolites (*i.e.* common adducts). For each bucket, the feature with the highest intensity was used for quantification based on the peak height in MS mode and only these features were included in the final dataset. Features in the retention time (RT) range 1.25 – 29 min (*i.e.* excluding the injection peak) were aligned across samples (ProfileAnalysis v. 2.3, Bruker Daltonics), allowing RT shifts of 0.1 min and m/z shifts of 6 mDa. Only features whose mean intensity was at least 50 times higher than in blanks and which occurred in at least two samples were retained in the dataset. For samples harvested in 2022, processing of the LC-MS data were done following the same steps and using similar parameters on the T-ReX 3D algorithm of MetaboScape (v. 2021b, Bruker Daltonics). Settings included the presence of features in minimum 3 samples, correlation coefficient threshold (ESI correlation) 0.8, intensity (peak height) threshold 1,000, minimum peak length 11. For both tables, features were normalised by dividing the peak heights by the height of the hydrocortisone [M+HCOOH-H]^-^ ion (chemical standard) and feature intensities were divided by the sample dry weights. GC-MS and LC-MS datasets were normalised by median normalisation, cube-root transformation and Pareto scaling before statistical analyses. Data normality was checked in MetaboAnalyst (v. 5.0)^55^. The non-normalised datasets as well as feature and sample metadata are available as supplemental data (Tables S1, S2, S8 and S9).

### Generalised multilinear models (GLMs)

To test the capacity of terpenoid profiles and metabolic fingerprints to predict the chemotype and maternal genotype, GLMs were conducted in R (v. 4.2.1) using the *glmnet* package^56^, as previously described^57^. Briefly, multinomial models were developed by testing a thousand penalty values of elastic net (ranging from 0 to 1) for variable selection. Stratified sampling was used to uniformly divide the sample set into training (60%), validation (20%) and testing (20%) sets. To cope with the random partitioning, 500 models were performed for each test. Cross-validation was applied to limit overfitting and prediction accuracy was defined on the real predictions performed on the test set. To statistically validate the models, the likelihood of spurious predictions was estimated using 500 permuted datasets, in which either chemotypes or maternal genotypes were randomly assigned to samples. Variable selection was performed using variable occurrence in the 500 models^57^ to select the best markers (*i.e.* features allowing for significant discrimination between classes). Model performance (*i.e.* prediction accuracy) was compared using Student t-tests. Finally, biological validation was performed using the complementary validation set. The equation of the model was calculated using the initial dataset (plants collected in the field in June 2021) and directly applied to the samples of the complementary set (collected in October 2022) to predict the chemotype or maternal genotype of these samples. The likelihood of spurious prediction was again defined using 500 permuted datasets.

### Multivariate statistical analyses

Principal component analyses (PCA) and variation partitioning were performed in R (v.4.2.1) using *FactoMineR* and *vegan* packages, respectively^58,59^. Tukey’s tests were performed in MetaboAnalyst (v5.0)^55^ to extract significant features (i < 0.05, FDR correction). Partial least square discriminant analyses were performed in MetaboAnalyst. To compare model performance, Tukey’s tests were done using the *agricolae* package. Box-whisker plots, heatmaps (Pearson correlation, Ward algorithm) and upset plots were created using *ggplot2, ggpubr, Hmisc, pheatmap* and *UpSetR* packages^60–65^, respectively.

### Annotation of LC-MS features

Putative molecular formulas of the best LC-MS markers were defined using SmartFormula and SmartFormula 3D in MetaboScape (v. 2021b) including N, O, C, H, S, Cl and P elements. The most likely metabolic candidates were selected based on the *m/z* deviation (Δppm) and were screened on chemical databases such as ChEBI, DNP (http://dnp.chemnetbase.com) and KNApSAcK^66,67^. When available, MS/MS spectra were compared to MS/MS spectra from MassBank to improve annotation confidence^68^. In addition, an in-house library was used to compare retention time, MS and MS/MS spectra. Confidence in the annotation level was defined according to the metabolomics standards initiative (MSI) confidence level^69^ (Table S5). For MSI 4 level (lower confidence level), the putative chemical class was assigned according to the most represented chemical class (*i.e.* the most widely represented chemical class for a given molecular formula) and based on the literature on tansy. For this confidence level, a putative compound was assigned as a potential example. As most of the putative annotations did not reach the MSI 2 level, the analysis of the results was mainly performed at the chemical class level. Biochemical pathways and putative chemical classes were inferred using KEGG identifiers^70^ and Classyfier (http://classyfire.wishartlab.com).

## Acknowledgements

We thank Tanja Bloss, Lukas Brokate, Stephanie Champion and Birte Wolf for assistance in the field and analytical work. We also thank Sylvain Prigent for his advice on the GLM-based modelling approach. The work was funded by the German Research Foundation (DFG), project MU 1829/28-1, as part of the Research Unit (RU) FOR 3000.

## Author contributions

TD and CM conceived the project. DZ, RJ and EJE participated in fieldwork and sampling. Terpenoid analyses were supervised and conducted by DZ, TTNN, LK and EJE; RS and CM performed the LC-MS measurements. Repeated metabolomics and bioinformatics analyses were conducted by TD, RS and CM. TD and CM wrote the manuscript with feedback from all co-authors.

## Additional information

We declare no potential conflict of interest.

## Data Availability

All data and metadata are available in supplemental tables and were deposited online ().

## Supplemental materials

**Fig. S1 | Effect of growing conditions on major terpenoid profile.**

**Fig. S2 | Distribution of the best chemotype markers between chemotypes.**

**Fig. S3 | Correlation between LC-MS and GC-MS predictors.**

**Fig. S4 | Effect of the maternal genotype on GC-MS and LC-MS data.**

**Fig. S5 | Maternal genotype effects on terpenoid profiles within chemotypes.**

**Fig. S6 | Maternal genotype effects on metabolic fingerprints within chemotypes.**

**Table S1. GC-MS data.**

**Table S2. LC-MS data of samples harvested in June 2021.**

**Table S3. Presence of the metabolic features in the chemotypes.**

**Table S4. Ranking of LC-MS features predicting chemotypes.**

**Table S5. Annotation of the best LC-MS predictors.**

**Table S6. Ranking of LC-MS features predicting maternal genotypes.**

**Table S7. Sample metadata.**

**Table S8. LC-MS data of samples harvested in October 2022 (complementary set).**

**Table S9. GC-MS metadata.**

## References

1. Holopainen, M., Hiltunen, R. & von Schantz, M. A Study on *Tansy Chemotypes*. Planta Med 53, 284–287 (1987).

2. Firn, R. D. & Jones, C. G. Natural products ? A simple model to explain chemical diversity. Natural Product Reports 20, 382 (2003).

3. Scossa, F. & Fernie, A. R. The evolution of metabolism: How to test evolutionary hypotheses at the genomic level. Computational and Structural Biotechnology Journal 18, 482–500 (2020).

4. Moore, B. D., Andrew, R. L., Külheim, C. & Foley, W. J. Explaining intraspecific diversity in plant secondary metabolites in an ecological context. New Phytologist 201, 733–750 (2014).

5. Weng, J.-K., Lynch, J. H., Matos, J. O. & Dudareva, N. Adaptive mechanisms of plant specialized metabolism connecting chemistry to function. Nature Chemical Biology 17, 1037–1045 (2021).

6. Kleine, S. & Müller, C. Intraspecific plant chemical diversity and its relation to herbivory. Oecologia 166, 175–186 (2011).

7. Moritz, F., Kaling, M., Schnitzler, J.-P. & Schmitt-Kopplin, P. Characterization of poplar metabotypes via mass difference enrichment analysis: mass difference network analysis in poplar. Plant, Cell & Environment 40, 1057–1073 (2017).

8. Clancy, M. V. et al. Metabotype variation in a field population of tansy plants influences aphid host selection: Plant chemical diversity in a plant-aphid system. Plant, Cell and Environment 41, 2791–2805 (2018).

9. Keskitalo, M., Pehu, E. & Simon, J. E. Variation in volatile compounds from tansy (*Tanacetum vulgare L.)*related to genetic and morphological differences of genotypes. Biochemical Systematics and Ecology 29, 267–285 (2001).

10. van Leur, H., Raaijmakers, C. E. & van Dam, N. M. A heritable glucosinolate polymorphism within natural populations of *Barbarea vulgaris*. Phytochemistry 67, 1214–1223 (2006).

11. Castells, E., Mulder, P. P. J. & Pérez-Trujillo, M. Diversity of pyrrolizidine alkaloids in native and invasive *Senecio pterophorus* (Asteraceae): implications for toxicity. Phytochemistry 108, 137–146 (2014).

12. Eilers, E. J., Kleine, S., Eckert, S., Waldherr, S. & Müller, C. Flower production, headspace volatiles, pollen nutrients, and florivory in *Tanacetum vulgare* chemotypes. Frontiers in Plant Science 11, 611877 (2021).

13. Calf, O. W. et al. Gastropods and insects prefer different *Solanum dulcamara* chemotypes. Journal of Chemical Ecology 45, 146–161 (2019).

14. Züst, T. et al. Independent evolution of ancestral and novel defenses in a genus of toxic plants (*Erysimum*, Brassicaceae). eLife 9, e51712 (2020).

15. Kazemi-Dinan, A., Sauer, J., Stein, R. J., Krämer, U. & Müller, C. Is there a trade-off between glucosinolate-based organic and inorganic defences in a metal hyperaccumulator in the field? Oecologia 178, 369–378 (2015).

16. Tewes, L. J., Michling, F., Koch, M. A. & Müller, C. Intracontinental plant invader shows matching genetic and chemical profiles and might benefit from high defence variation within populations. Journal of Ecology 106, 714–726 (2018).

17. Fortuna, T. M. et al. Variation in plant defences among populations of a range-expanding plant: consequences for trophic interactions. New Phytologist 204, 989–999 (2014).

18. Clancy, M. V. et al. Terpene chemotypes in *Gossypium hirsutum* (wild cotton) from the Yucatan Peninsula, Mexico. Phytochemistry 205, 113454 (2023).

19. Holopainen, M., Hiltunen, R., Lokki, J., Forsén, K. & Schantz, M. V. Model for the genetic control of thujone, sabinene and umbellulone in tansy (*Tanacetum vulgare L.)*. Hereditas 106, 205–208 (1997).

20. Schuman, M. C., van Dam, N. M., Beran, F. & Harpole, W. S. How does plant chemical diversity contribute to biodiversity at higher trophic levels? Current Opinion in Insect Science 14, 46–55 (2016).

21. Kleine, S. & Müller, C. Drought stress and leaf herbivory affect root terpenoid concentrations and growth of *Tanacetum vulgare*. Journal of Chemical Ecology 40, 1115–1125 (2014).

22. Jakobs, R., Schweiger, R. & Müller, C. Aphid infestation leads to plant part-specific changes in phloem sap chemistry, which may indicate niche construction. New Phytologist 221, 503–514 (2019).

23. Lokki, J., Sorsa, M., Forsén, K. & Schantz, M. V. Genetics of monoterpenes in *Chrysanthemum vulgare:* I. Genetic control and inheritance of some of the most common chemotypes. Hereditas 74, 225–232 (1973).

24. Thompson, J. D., Chalchat, J.-C., Michet, A., Linhart, Y. B. & Ehlers, B. Qualitative and quantitative variation in monoterpene co-occurrence and composition in the essential oil of *Thymus vulgaris* chemotypes. Journal of Chemical Ecology 29, 859–880 (2003).

25. Johnson, R. H., Hull-Sanders, H. M. & Meyer, G. A. Comparison of foliar terpenes between native and invasive *Solidago gigantea*. Biochemical Systematics and Ecology 35, 821–830 (2007).

26. Booth, J. K. & Bohlmann, J. Terpenes in *Cannabis sativa* – from plant genome to humans. Plant Science 284, 67–72 (2019).

27. Wolf, V. C., Gassmann, A., Clasen, B. M., Smith, A. G. & Müller, C. Genetic and chemical variation of *Tanacetum vulgare* in plants of native and invasive origin. Biological Control 61, 240–245 (2012).

28. Linhart, Y. B., Keefover-Ring, K., Mooney, K. A., Breland, B. & Thompson, J. D. A chemical polymorphism in a multitrophic setting: Thyme monoterpene composition and food web structure. The American Naturalist 166, 517–529 (2005).

29. Hao, D.-C. et al. The genus Chrysanthemum: Phylogeny, biodiversity, phytometabolites, and chemodiversity. Frontiers in Plant Science 13, 973197 (2022).

30. Ak, G. et al. *Tanacetum vulgare L.* (Tansy) as an effective bioresource with promising pharmacological effects from natural arsenal. Food and Chemical Toxicology 153, 112268 (2021).

31. Goławska, S. & Łukasik, I. Antifeedant activity of luteolin and genistein against the pea aphid, *Acyrthosiphon pisum*. Journal of Pest Science 85, 443–450 (2012).

32. Roddy, A. B. et al. Towards the flower economics spectrum. New Phytologist 229, 665–672 (2021).

33. Kleine, S. & Müller, C. Differences in shoot and root terpenoid profiles and plant responses to fertilisation in *Tanacetum vulgare*. Phytochemistry 96, 123–131 (2013).

34. Fang, C., Fernie, A. R. & Luo, J. Exploring the diversity of plant metabolism. Trends in Plant Science 24, 83–98 (2019).

35. Yonekura-Sakakibara, K., Higashi, Y. & Nakabayashi, R. The origin and evolution of plant flavonoid metabolism. Frontiers in Plant Science 10, 943 (2019).

36. Yoshikuni, Y., Ferrin, T. E. & Keasling, J. D. Designed divergent evolution of enzyme function. Nature 440, 1078–1082 (2006).

37. Wittmann, M. J. & Fukami, T. Eco-evolutionary buffering: rapid evolution facilitates regional species coexistence despite local priority effects. The American Naturalist 191, E171–E184 (2018).

38. Tholl, D. Terpene synthases and the regulation, diversity and biological roles of terpene metabolism. Current Opinion in Plant Biology 9, 297–304 (2006).

39. Gershenzon, J. & Dudareva, N. The function of terpene natural products in the natural world. Nature Chemical Biology 3, 408–414 (2007).

40. Pichersky, E. & Gershenzon, J. The formation and function of plant volatiles: perfumes for pollinator attraction and defense. Current Opinion in Plant Biology 5, 237–243 (2002).

41. Junker, R. R., Gershenzon, J. & Unsicker, S. B. Floral odor bouquet loses its ant repellent properties after inhibition of terpene biosynthesis. Journal of Chemical Ecology 37, 1323–1331 (2011).

42. Boeckler, G. A., Gershenzon, J. & Unsicker, S. B. Phenolic glycosides of the Salicaceae and their role as anti-herbivore defenses. Phytochemistry 72, 1497–1509 (2011).

43. Barbehenn, R. V. & Peter Constabel, C. Tannins in plant–herbivore interactions. Phytochemistry 72, 1551–1565 (2011).

44. Devrnja, N. et al. Comparative studies on the antimicrobial and cytotoxic activities of *Tanacetum vulgare L.* essential oil and methanol extracts. South African Journal of Botany 111, 212–221 (2017).

45. Acimovic, M. & Puvača, N. *Tanacetum vulgare L.* - A Systematic Review. Journal of Agronomy, Technology and Engineering Management 3, 416–422 (2020).

46. Bais, H. P., Weir, T. L., Perry, L. G., Gilroy, S. & Vivanco, J. M. The role of root exudates in rhizosphere interactions with plants and other organisms. Annual Review of Plant Biology 57, 233–266 (2006).

47. Rasmann, S., Bauerle, T. L., Poveda, K. & Vannette, R. Predicting root defence against herbivores during succession: Root defence against herbivores. Functional Ecology 25, 368–379 (2011).

48. Singh, P., Arif, Y., Bajguz, A. & Hayat, S. The role of quercetin in plants. Plant Physiology and Biochemistry 166, 10–19 (2021).

49. Junker, R. R. & Blüthgen, N. Floral scents repel facultative flower visitors, but attract obligate ones. Annals of Botany 105, 777–782 (2010).

50. Upchurch, R. G. Fatty acid unsaturation, mobilization, and regulation in the response of plants to stress. Biotechnology Letters 30, 967–977 (2008).

51. Desjardins, A. E. Natural product chemistry meets genetics: When Is a genotype a chemotype? Journal of Agriculture and Food Chemistry 56, 7587–7592 (2008).

52. Ziaja, D. & Müller, C. Intraspecific chemodiversity provides plant individual- and neighbourhood-mediated associational resistance towards aphids. http://biorxiv.org/lookup/doi/10.1101/2022.12.21.521353 (2022) doi:10.1101/2022.12.21.521353.

53. Schweiger, R., Castells, E., Da Sois, L., Martínez-Vilalta, J. & Müller, C. Highly species-specific foliar metabolomes of diverse woody species and relationships with the leaf economics spectrum. Cells 10, 644 (2021).

54. van Den Dool, H. & Dec. Kratz, P. A generalization of the retention index system including linear temperature programmed gas—liquid partition chromatography. Journal of Chromatography A 11, 463–471 (1963).

55. Pang, Z. et al. MetaboAnalyst 5.0: narrowing the gap between raw spectra and functional insights. Nucleic Acids Research 49, W388–W396 (2021).

56. Friedman, J., Hastie, T. & Tibshirani, R. Regularization paths for generalized linear models via coordinate descent. Journal of Statistical Software 33, 1–22 (2010).

57. Dussarrat, T. et al. Predictive metabolomics of multiple Atacama plant species unveils a core set of generic metabolites for extreme climate resilience. New Phytologist nph.18095 (2022) doi:10.1111/nph.18095.

58. Lê, S., Josse, J. & Husson, F. FactoMineR: A package for multivariate analysis. Journal of Statistical Software 25, 1–18 (2008).

59. Oksanen, J. et al. vegan: Community Ecology Package. (2022).

60. Wickham, H. ggplot2: Elegant Graphics for Data Analysis. (Springer-Verlag New York, 2016).

61. Kolde, R. pheatmap: Pretty Heatmaps. (2019).

62. Jr, F. E. H., Dupont, with contributions from C. & others, many. Hmisc: Harrell Miscellaneous. (2020).

63. Kassambara, A. ggpubr: ‘ggplot2’ Based Publication Ready Plots. (2020).

64. Mendiburu, F. de. agricolae: Statistical Procedures for Agricultural Research. (2021).

65. Conway, J. R., Lex, A. & Gehlenborg, N. UpSetR: an R package for the visualization of intersecting sets and their properties. Bioinformatics 33, 2938–2940 (2017).

66. Afendi, F. M. et al. KNApSAcK family databases: integrated metabolite-plant species databases for multifaceted plant research. Plant Cell Physiol. 53, e1 (2012).

67. Hastings, J. et al. ChEBI in 2016: Improved services and an expanding collection of metabolites. Nucleic Acids Res 44, D1214–D1219 (2016).

68. Horai, H. et al. MassBank: a public repository for sharing mass spectral data for life sciences. J. Mass Spectrom. 45, 703–714 (2010).

69. Sumner, L. W. et al. Proposed quantitative and alphanumeric metabolite identification metrics. Metabolomics 10, 1047–1049 (2014).

70. Kanehisa, M. et al. KEGG for linking genomes to life and the environment. Nucleic Acids Research 36, D480–D484 (2007).

