## Supplemental Figure S1 for "Influences of chemotype and parental genotype on metabolic fingerprints of tansy plants uncovered by predictive metabolomics"

### Slide 1
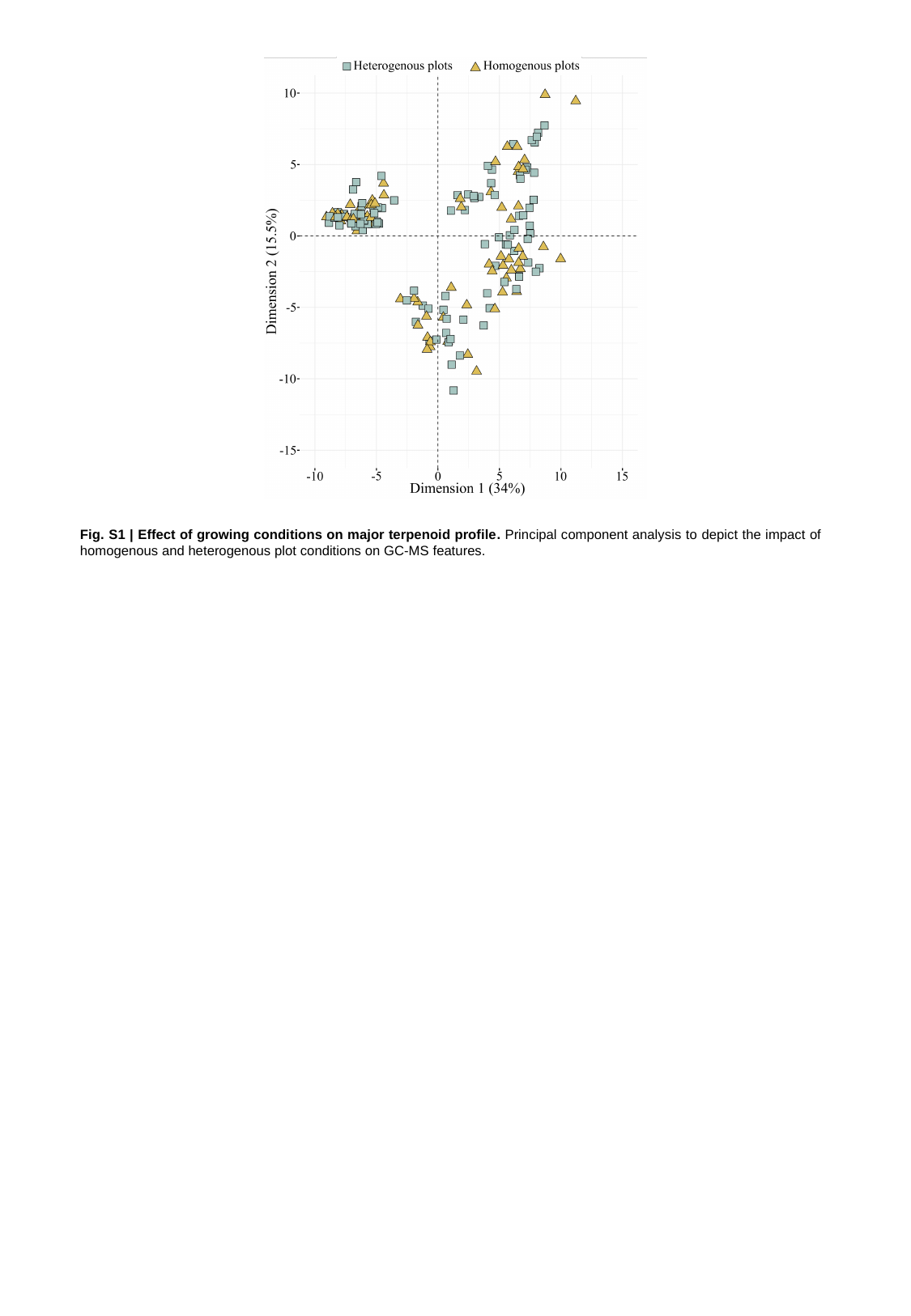

Fig. S1 | Effect of growing conditions on major terpenoid profile. Principal component analysis to depict the impact of homogenous and heterogenous plot conditions on GC-MS features.
