## Supplemental Figure S2 for "Influences of chemotype and parental genotype on metabolic fingerprints of tansy plants uncovered by predictive metabolomics"

### Slide 1
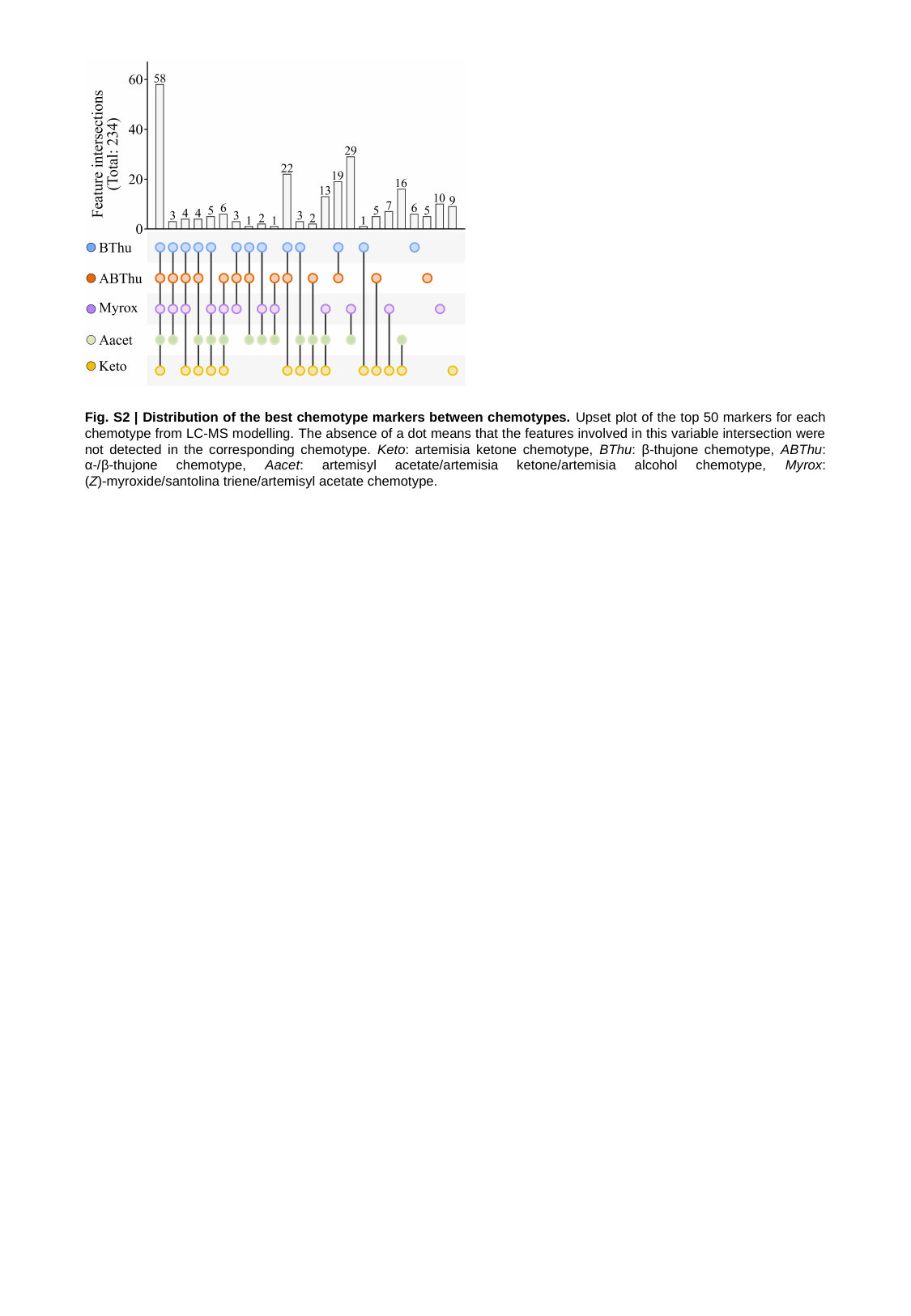

Fig. S2 | Distribution of the best chemotype markers between chemotypes. Upset plot of the top 50 markers for each chemotype from LC-MS modelling. The absence of a dot means that the features involved in this variable intersection were not detected in the corresponding chemotype. Keto: artemisia ketone chemotype, BThu: β-thujone chemotype, ABThu: α-/β-thujone chemotype, Aacet: artemisyl acetate/artemisia ketone/artemisia alcohol chemotype, Myrox: (Z)-myroxide/santolina triene/artemisyl acetate chemotype.
