## Supplemental Figure S3 for "Influences of chemotype and parental genotype on metabolic fingerprints of tansy plants uncovered by predictive metabolomics"

### Slide 1
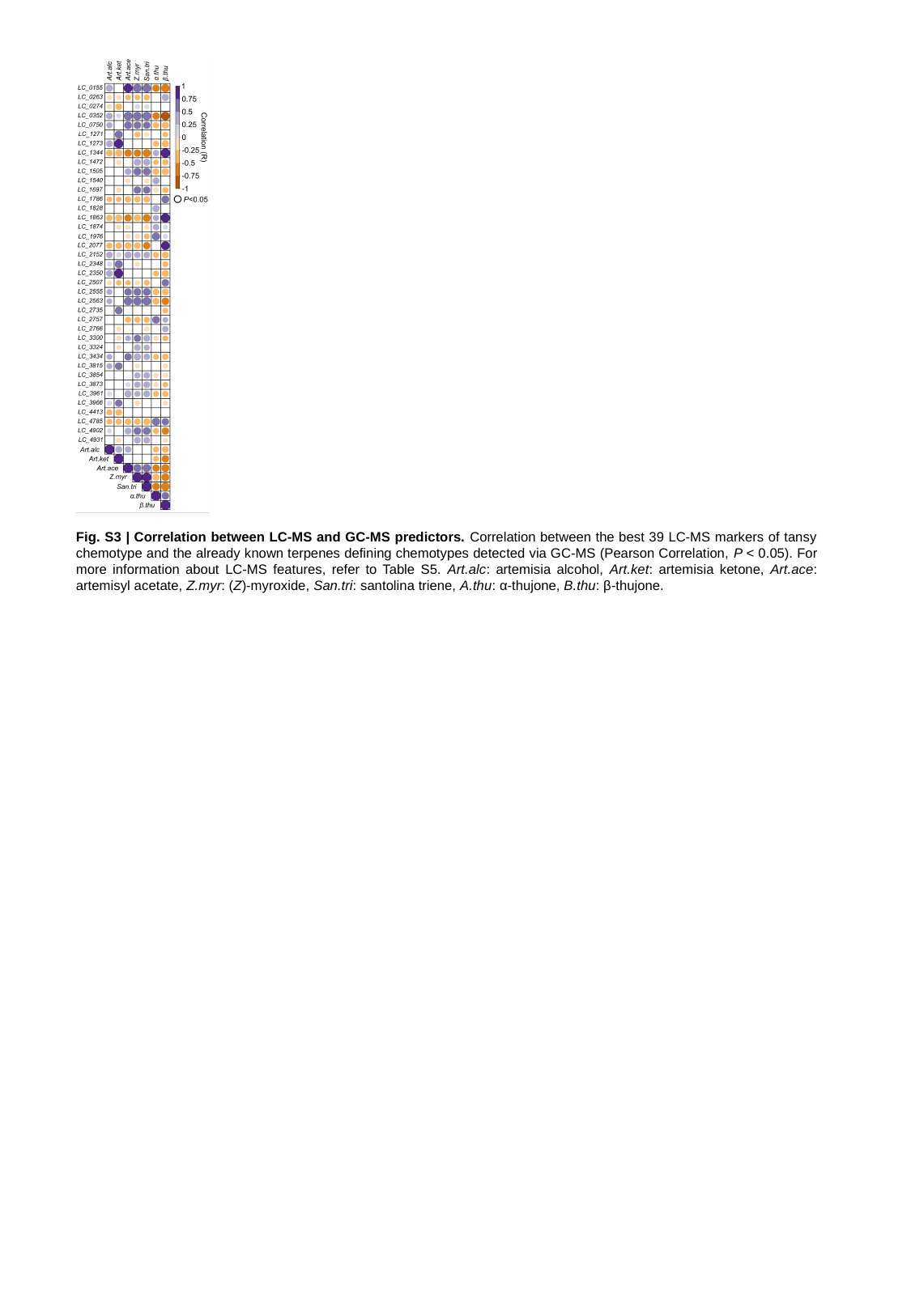

Fig. S3 | Correlation between LC-MS and GC-MS predictors. Correlation between the best 39 LC-MS markers of tansy chemotype and the already known terpenes defining chemotypes detected via GC-MS (Pearson Correlation, P < 0.05). For more information about LC-MS features, refer to Table S5. Art.alc: artemisia alcohol, Art.ket: artemisia ketone, Art.ace: artemisyl acetate, Z.myr: (Z)-myroxide, San.tri: santolina triene, A.thu: α-thujone, B.thu: β-thujone.
