## Supplemental Figure S4 for "Influences of chemotype and parental genotype on metabolic fingerprints of tansy plants uncovered by predictive metabolomics"

### Slide 1
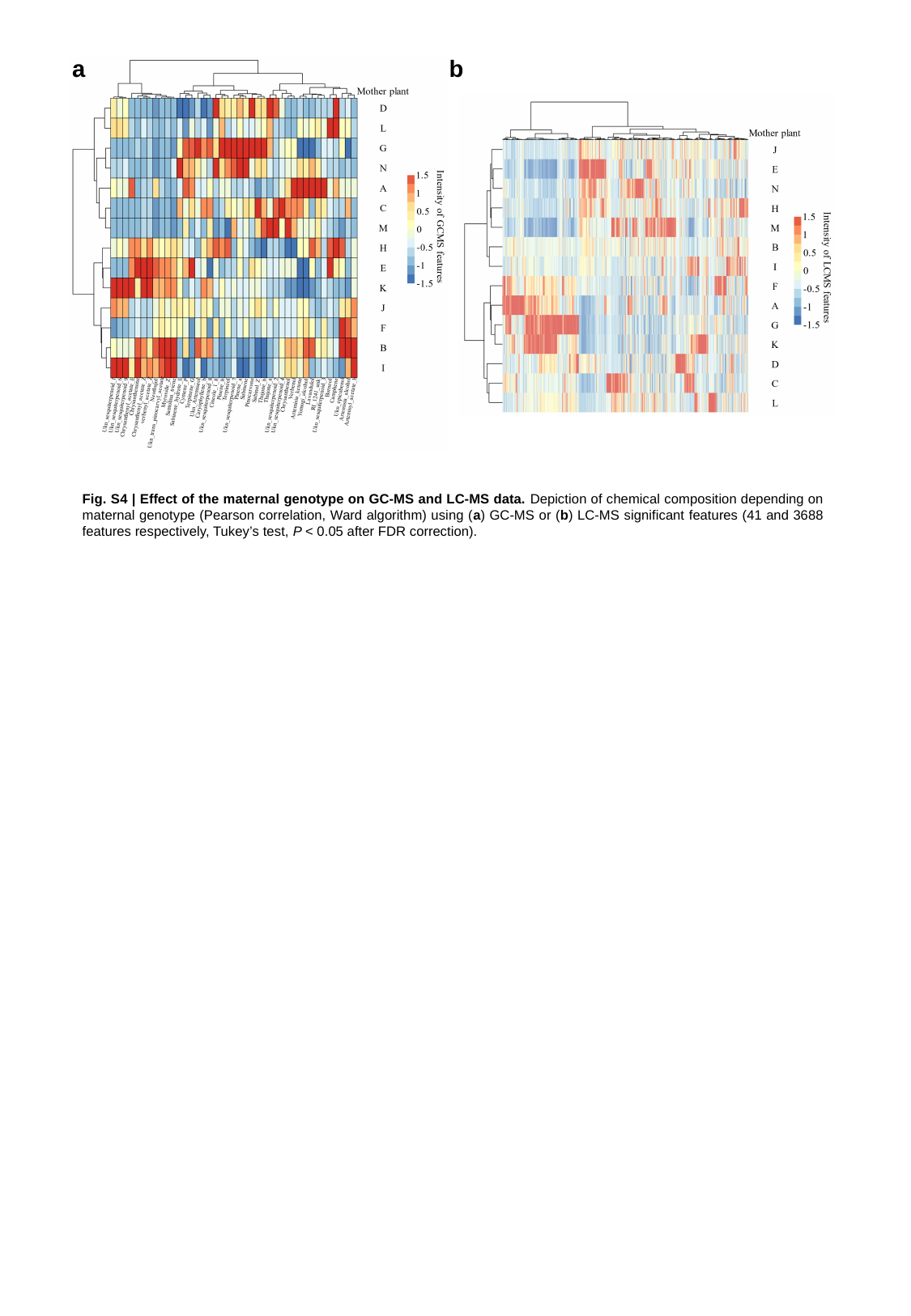

a
b
Fig. S4 | Effect of the maternal genotype on GC-MS and LC-MS data. Depiction of chemical composition depending on maternal genotype (Pearson correlation, Ward algorithm) using (a) GC-MS or (b) LC-MS significant features (41 and 3688 features respectively, Tukey’s test, P < 0.05 after FDR correction).
