## Supplemental Figure S6 for "Influences of chemotype and parental genotype on metabolic fingerprints of tansy plants uncovered by predictive metabolomics"

### Slide 1
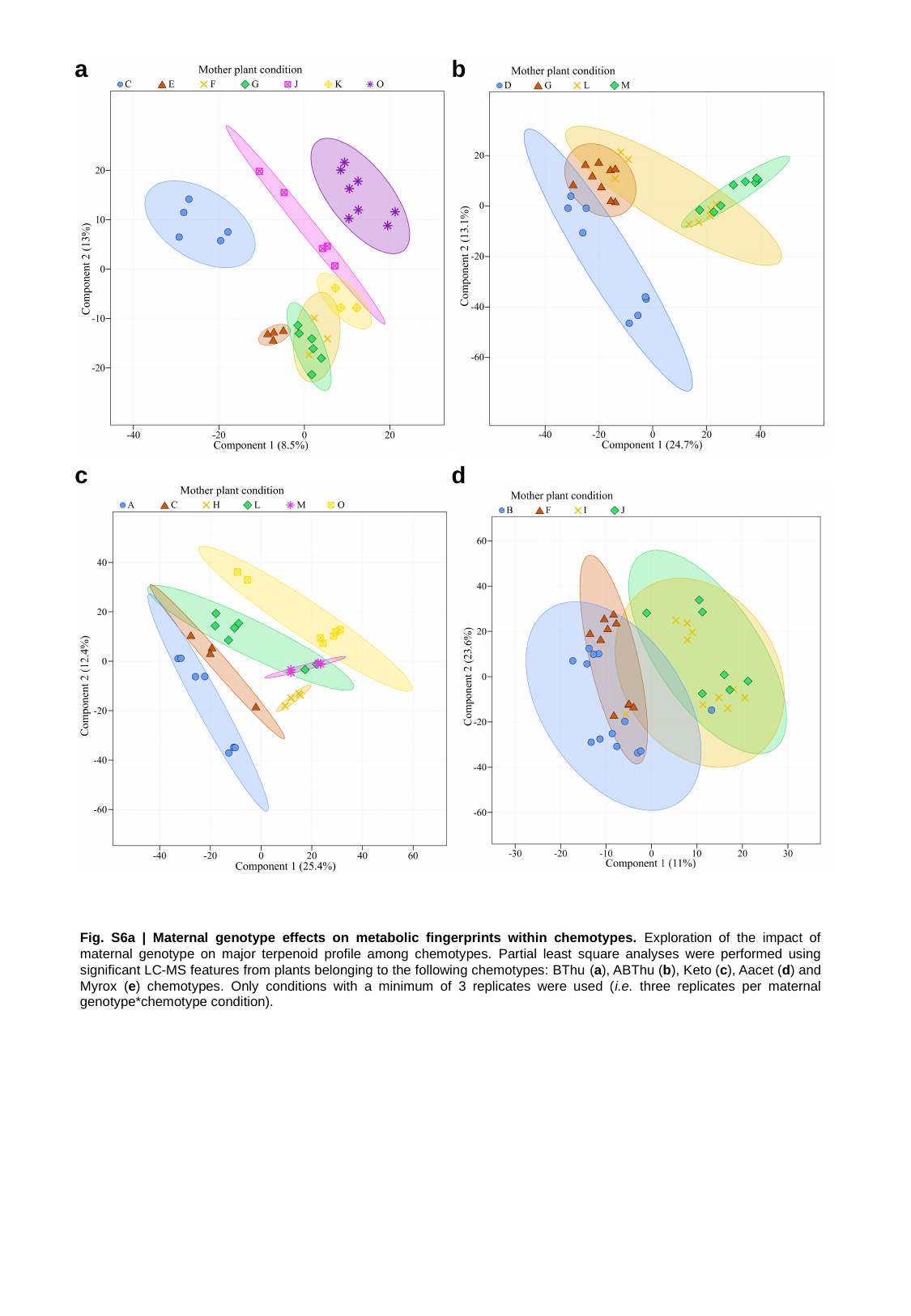

a
b
c
d
Fig. S6a | Maternal genotype effects on metabolic fingerprints within chemotypes. Exploration of the impact of maternal genotype on major terpenoid profile among chemotypes. Partial least square analyses were performed using significant LC-MS features from plants belonging to the following chemotypes: BThu (a), ABThu (b), Keto (c), Aacet (d) and Myrox (e) chemotypes. Only conditions with a minimum of 3 replicates were used (i.e. three replicates per maternal genotype*chemotype condition).

### Slide 2
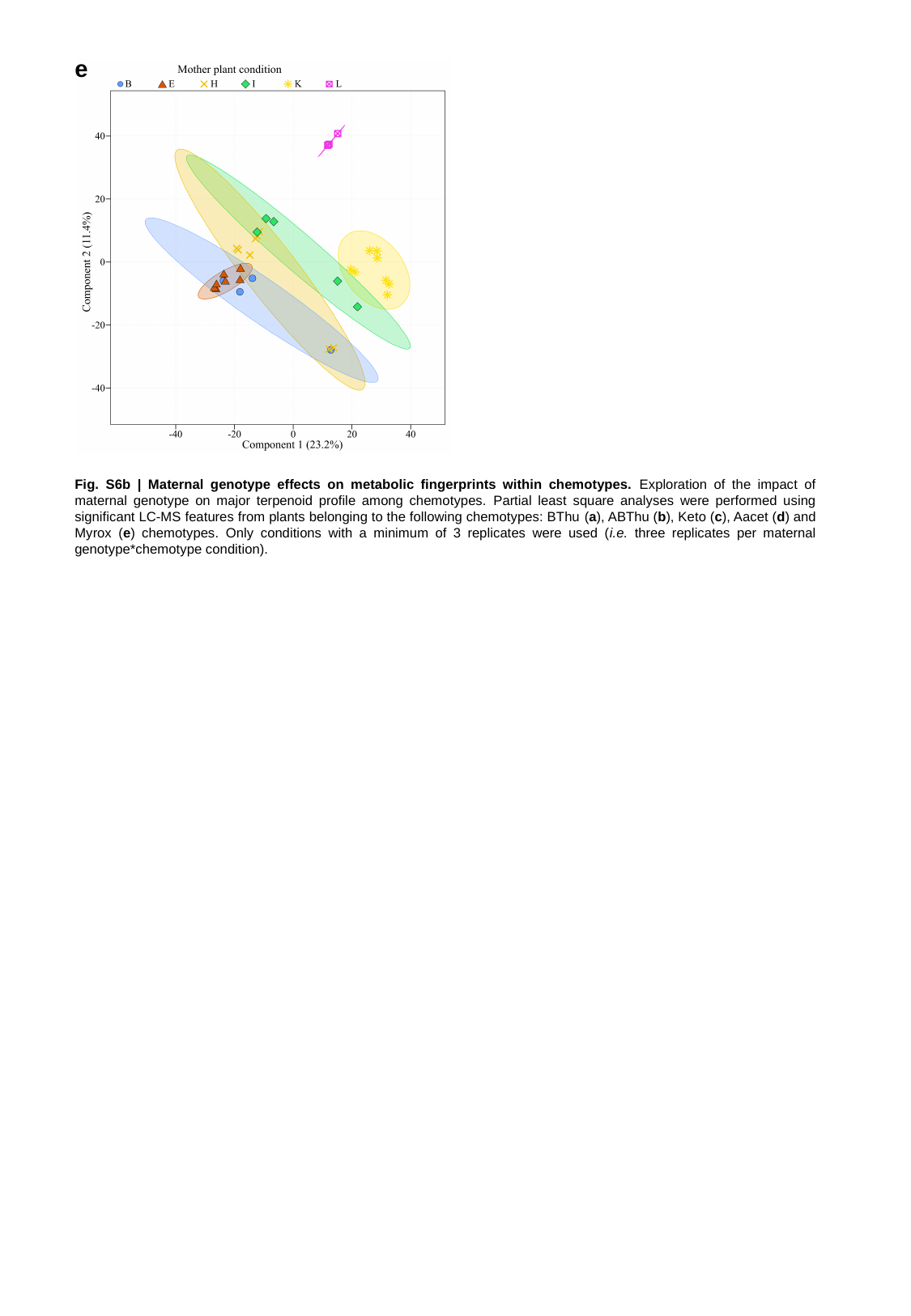

e
Fig. S6b | Maternal genotype effects on metabolic fingerprints within chemotypes. Exploration of the impact of maternal genotype on major terpenoid profile among chemotypes. Partial least square analyses were performed using significant LC-MS features from plants belonging to the following chemotypes: BThu (a), ABThu (b), Keto (c), Aacet (d) and Myrox (e) chemotypes. Only conditions with a minimum of 3 replicates were used (i.e. three replicates per maternal genotype*chemotype condition).
